# Finger representations in primary somatosensory cortex are modulated by a vibrotactile working memory task

**DOI:** 10.1101/2021.10.29.466459

**Authors:** Finn Rabe, Sanne Kikkert, Nicole Wenderoth

## Abstract

It is well-established that several cortical areas represent vibrotactile stimuli in somatotopic maps. However, whether such somatotopic representations remain active during the delay period of working memory (WM) tasks, i.e. in the absence of any tactile stimulation, is unknown. In our experiment, participants had to compare two tactile stimuli with different vibration frequencies that were separated by a delay period (memory condition) or they were exposed to identical stimuli but did not have to solve a WM task (no memory condition). Importantly, both vibrotactile stimuli were either applied to the right index or little finger. Analyzing the delay period, we identified a well-known fronto-parietal network of brain regions involved in WM but we did not find WM specific activity in S1. However, using multi-voxel pattern analysis (MVPA) and representational similarity analysis (RSA), we found that S1 finger representations were more dissimilar during the delay period of the WM condition than during the control condition. These results indicate that WM processes modulate the representational geometry of S1 suggesting that some aspects of the tactile WM content are represented in a somatotopic fashion.

**HIGHLIGHTS:**

- Multivariate approaches were used to identify finger specific representational changes during vibrotactile frequency discrimination.
- Vibrotactile working memory modulates somatotopic finger representations in contralateral S1 during the delay period, i.e. in the absence of any tactile stimuli

## 1. INTRODUCTION

Topographic representations such as the somatotopic map in the somatosensory cortex have been shown to be ubiquitous in the cerebral cortex of mammals. They consist of orderly representations of receptor surfaces on different body parts (Kaas, 1993, 1997; Penfield &Boldrey, 1937; Silver &Kastner, 2009). These somatotopic maps are incredibly specific to the point where the sensation of each finger can be assigned to its own cortical region, so called finger representations (Besle et al., 2013; Martuzzi et al., 2014; Sanchez Panchuelo et al., 2018; Sanders et al., 2019).

Importantly, finger-specific somatosensory representations are not exclusively activated by tactile or proprioceptive stimulation, but they are also modulated through other mechanisms like (i) *attempted movement*s which do not produce overt motor output or the associated tactile or proprioceptive feedback (Ariani et al., 2021; Guan et al., 2021; Kikkert et al., 2021) (ii) *observed* (Kuehn et al., 2018) or *actively imagined* touch (Schmidt &Blankenburg, 2019), (iii) or directing *attention* to a specific finger (Puckett et al., 2017).

Another mechanism that may modulate S1 activity is tactile working memory (WM) i.e. when somatosensory stimuli have to be kept in memory beyond the actual stimulation period for subsequent decision making (Christophel et al., 2017). Support for S1 involvement in tactile WM stems from neurophysiological research in non-human primates. Single-unit activity was recorded from the S1 hand area while subjects had to match an object with a specific surface to a previously presented surface stimulus (Y. D. Zhou &Fuster, 1996; Y.-D. Zhou &Fuster, 2000). The authors observed cells that were not only active while the tactile stimulus was present, but also sustained their firing during the WM delay period. This suggests that primary sensory cortices could serve as a memory buffer for stimulus information (D’Esposito &Postle, 2015), an idea that has been conceptualized as the ‘sensory recruitment’ model of WM (Katus et al., 2015; Pasternak &Greenlee, 2005). According to this model, WM is maintained in those brain regions that are involved in encoding sensory stimuli.

However, human neuroimaging studies on vibrotactile WM have revealed mixed results as to whether S1 is activated during the delay period of information storage: Several studies have shown that the average activity level of S1 (as detected by a “mass-univariate” statistical approach) is not significantly larger during the WM delay period than during a control condition (for reviews see (Christophel et al., 2017). Dynamical causal modelling (DCM) estimates of interactions between brain regions suggested that during vibrotactile frequency discrimination somatosensory stimuli are serially processed from S1 to secondary somatosensory cortex (S2;(Kalberlah et al., 2013), which matches findings in non-human primates (for review (Romo &Rossi-Pool, 2020).

Other studies, by contrast, have employed multi-voxel pattern analysis (MVPA) which can detect stimulus information in spatially distributed patterns of activation in a region of interest (ROI) (Weaverdyck et al., 2020). Neuroimaging studies utilizing MVPA found that features like vibratory frequencies were represented during the delay period, mainly in associative brain regions, i.e., posterior parietal and frontal regions (Schmidt et al., 2017; Wu et al., 2018). Interestingly, features of spatial layout stimuli could be decoded from S1 during the delay period (Schmidt &Blankenburg, 2018), even though such stimulus representations might fade out and in over time, especially when a temporal structure of the delay period can be anticipated (Rose et al., 2016). Similar to the aforementioned observations in non-human primates, during the delay period spatial layouts could be decoded from an area that usually represents the hand (Schmidt &Blankenburg, 2018). This area has been characterized by its fine-grained finger representations (Besle et al., 2013; Martuzzi et al., 2014; Sanchez Panchuelo et al., 2018; Sanders et al., 2019). It is, however, unknown whether somatotopic finger representations are modulated by a tactile WM task. To answer this question, we combined fMRI with multivariate analysis techniques to explore whether finger representations in contralateral S1 are selectively modulated when tactile stimuli are kept in WM. Participants were asked to perform a vibrotactile frequency discrimination task on the index or little finger while we collected fMRI data. We argue that above chance level classification accuracies and greater representational dissimilarities indicate that brain activation patterns representing single fingers in S1 are highly specific, in accordance with the idea of narrow tuning curves in the S1 neuronal population (Detorakis &Rougier, 2014). Lower classification accuracies and dissimilarities between fingers, by contrast, indicate that S1 activity is less finger specific suggesting that tuning curves are broader. Importantly, multivariate analysis can reveal representational information even if the average Blood-oxygen-level-dependent (BOLD) response does not surpass a certain statistical threshold in univariate analyses. We hypothesized that if WM selectively activates S1, then we will observe finger selective activation patterns during the WM delay period, reflected by greater classification accuracies and representational dissimilarity than during no memory delay periods.

## 2. MATERIAL AND METHODS

### 2.1. PARTICIPANTS

Thirty young healthy volunteers (19 females; mean age= 24.48, SEM= 0.44) participated in our study. Our sample size was comparable to those in previous reports on fMRI decoding of WM content using discrimination tasks (Ester et al., 2009; Schmidt et al., 2017). All participants were neurologically intact and reported to be right-handed. All of them gave written informed consent and the study protocol was approved by the local ethics committee (BASEC-Nr. 2018-01078). Three participants had to be excluded due to excessive head motion based on our criterion (see ‘Preprocessing of fMRI data’ section for more detail).

### 2.2. EXPERIMENTAL PROCEDURE AND TASKS

#### 2.2.1. TACTILE STIMULI

Vibrotactile stimuli (duration = 2s, sampling rate = 1kHz) were applied to the right index or right little finger using a MR-compatible piezoelectric device (PTS-T1, Dancer Design, UK). We selected these fingers as they have the largest inter-finger somatotopic distance (Besle et al., 2013; Ejaz et al., 2015; Kolasinski et al., 2016; Sanders et al., 2019), allowing us to robustly detect the modulation of somatotopic representations by the WM task. The one bin piezoelectric wafers were mounted to the fingertips using custom 3D-printed retainers that were fixed with a Velcro strap. Participants were asked to report any tingling sensation in case the retainer was mounted too tightly. The stimulation consisted of mechanical sinusoids that were transmitted from the testing computer to the piezoelectric device using a C Series Voltage Output Module (National Instruments) and the in-house NI-DAQmx driver.

#### 2.2.2. SENSORY DETECTION THRESHOLD ESTIMATION

To ensure similar task difficulty across runs of the main experiment (Harris et al., 2006), we determined the sensory detection threshold (SDT) for both fingers prior to starting the main experiment. SDT was defined as the stimulation intensity at which the participants detected the stimulus 50% of the time. We stimulated each finger only once per trial at base frequency (20 Hz) and participants were asked to press a button upon detection of a stimulation. To reliably estimate SDT, we applied a conventional Bayesian-based Quest procedure (QuestHandler in PsychoPy). After each detected or undetected stimulus the algorithm searched for the most probable psychometric function via maximum likelihood estimation over the course of 25 trials starting with a stimulation amplitude of 0.1 Volts (Watson &Pelli, 1983). The Weibull psychometric function was calculated using the following formula:

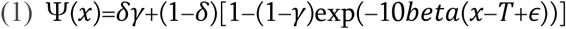

where χ is the stimulus intensity in Volts and *T* is the estimated sensory detection threshold. This procedure was performed prior to the first run. If the percentage of correctly discriminated memory trials in a run was below 60% or above 90%, then we redetermined the SDT using a shortened version of the Quest procedure. In such a case we started the Quest procedure with the previously determined stimulation intensity to reduce the number of iterations (new iterations = 7 trials). This procedure was applied to keep task difficulty at comparable levels throughout the experiment.

We analyzed changes in behavioral performance occurred across runs, potentially due to perceptual learning, cooling of the fingertips, or fatiguing effects using a repeated measures two-way ANOVA (2 fingers x 4 runs) (**Fig. 2B**).

#### 2.2.3. MAIN EXPERIMENTAL TASK

The main experimental task was generated using PsychoPy (Peirce et al., 2019). The experimental task consisted of memory and no memory trials. During a memory trial, participants performed a two-alternative forced choice (2AFC) discrimination task. Two vibrotactile stimuli were consecutively applied to the same finger (i.e., the index or the little finger), separated by a jittered 6-8s delay. We targeted cutaneous mechanoreceptors that respond to stimulations in the flutter range (Mountcastle et al., 1967). One of two stimuli vibrated at 20 Hz (2s duration at SDT intensity) while the vibration frequency of the other stimulus varied between 22, 24 or 26 Hz (same duration and intensity). Participants had to indicate by means of a button press whether the first or the second stimulation was higher in frequency (half of the participants) or whether the first or the second stimulation was lower in frequency (the other half of the participants), following previously published procedures (Pleger et al., 2006, 2008, 2009). Responses were recorded via index and middle finger button presses of the other (left) hand using a MR-compatible fiber optic device. We randomized the order of how the response options (f1 and f2) appeared on the screen on a trial-by-trail basis to prevent somatotopy-specific anticipatory motor activity. After a 3s response period participants received visual feedback (1s) indicating whether their response was correct (highlighted by green color) or incorrect (red; **Fig 1**). Participants were instructed to focus their gaze on the fixation cross in the middle of the screen during the complete trial. Vibrotactile stimuli trials targeted either the index or the little finger and which finger would be stimulated per trial was counterbalanced across each run.

**Fig 1.**
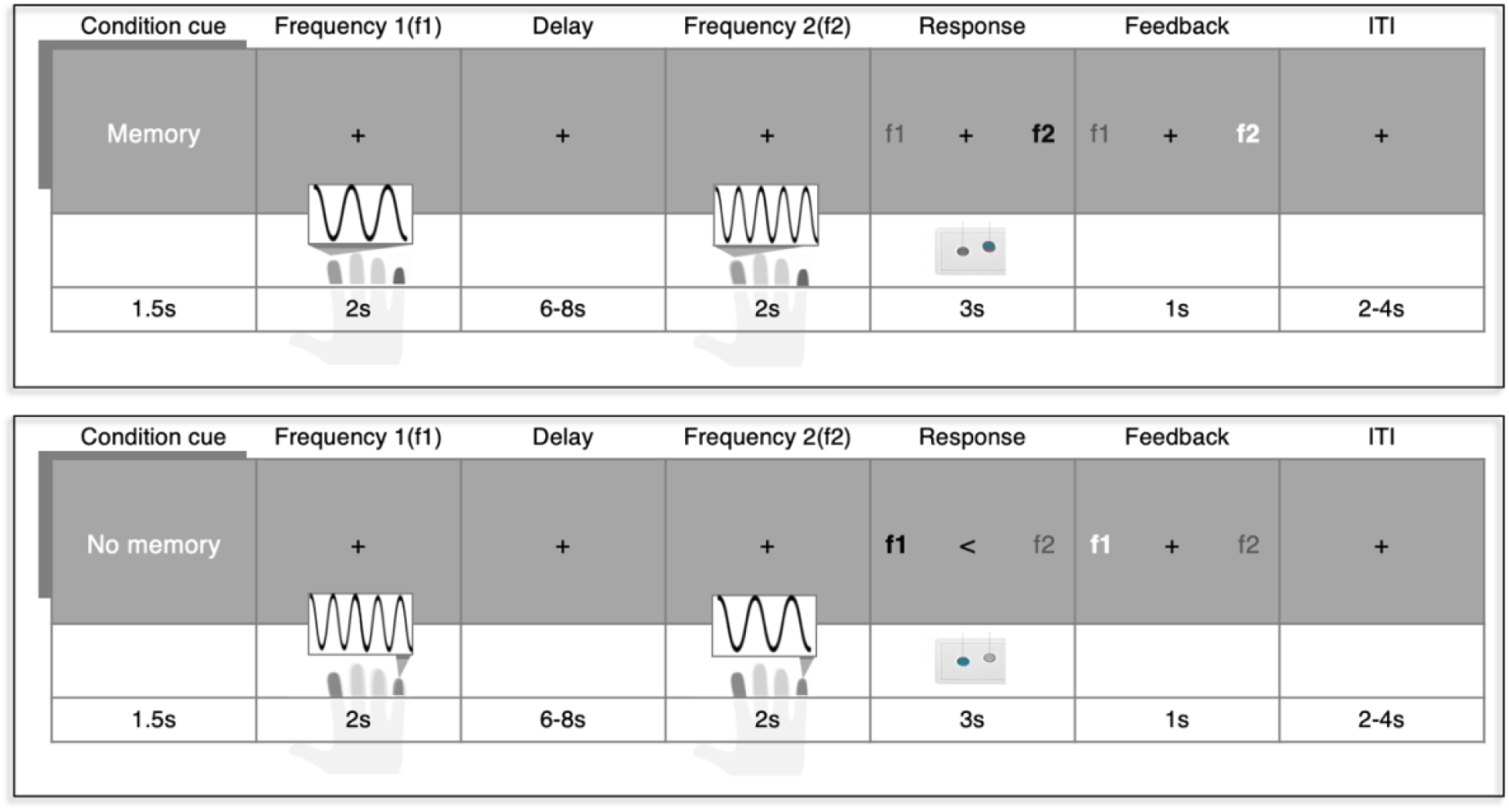
Vibrotactile frequency discrimination task. During memory trials (*top*) two vibrotactile stimuli that differed in frequency were consecutively applied to the same finger (in this example the index finger). Both stimulations were separated by a jittered delay period, during which participants had to keep the first stimulation frequency (f1) in memory in order to compare it to the frequency of the second stimulus (f2). During the 3s response period participants had to indicate which of the two stimulation frequencies (f1 and f2) was higher by means of a left hand button press. The mapping between the discrimination response and which button to press was indicated on the screen and randomized across trials. Subjects received feedback whether their response was correct or incorrect. The target finger (index or little finger) were intermixed within a run and the inter trial interval (ITI) was jittered between 2-4s. During no memory trials (*bottom*), vibrotactile stimulations and visual information remained the same. However, participants were instructed not to focus on the stimulation and also not to compare the vibrotactile frequencies. They simply had to press the button indicated by the arrow in the middle of the screen.

To disentangle WM processes from general responses to the stimuli, we also included no memory trials. During a no memory trial participants received the exact same vibratory stimulations as during memory trials, but they were instructed not to focus on the stimuli or on their vibration frequencies. During the response period subjects were informed by a visual cue (pointing arrow) which button to press. To ensure participants did not switch cognitive strategies, the indicated response was always contrary to the response that would be expected when correctly discriminating both frequencies. Memory and no memory trials conditions were separated in mini blocks of 4 trials. Participants were informed whether they had to perform the memory or no memory task by means of a visual cue (1.5s) at the beginning of each trial. Prior to the experiment, participants were familiarized with the memory and no memory tasks by completing 12 trials.

The order of stimulus sites (stimulated finger) was counterbalanced both within and across mini blocks. Stimulation frequencies were counterbalanced across the experiment. Each stimulus frequency was presented equally often in both memory and no memory condition. Jittered timings for Inter-stimulus-interval (ISI, 6-8s) and Inter-trial-interval (ITI, 2-4s) were randomly drawn from a uniform distribution. All participants completed 4 runs consisting of 48 trials each. Each run consisted of 6 memory and 6 no memory mini blocks in a counterbalanced order.

### 2.3. BEHAVIORAL ANALYSIS

We defined the discrimination accuracy per participant as the percentage of correctly discriminated trials separately for each condition. We expected that greater frequency differences would facilitate discrimination between both tactile vibrations while the stimulus site should have no effect. We therefore investigated whether behavioral performances differed across frequency differences and across fingers using a two-way repeated-measures ANOVA.

### 2.4. MRI DATA ACQUISITION

Functional as well as structural MRI images were acquired on a Philips Ingenia 3 Tesla MRI (Best, The Netherlands) using a 32-element head coil. fMRI data was collected using an echo-planar-imaging (EPI) sequence acquiring 36 transversal slices centred at the bicommissural line and with whole brain coverage, though excluding most of cerebellum (repetition time (TR): 2s, echo time (TE): 30ms, spatial resolution: 3mm^3^, FOV = 222 × 222mm^2^, 85° flip angle, slice orientation: transversal, SENSE factor (AP): 2, 472 functional volumes per run). Anatomical images were acquired during SDT estimation using a MPRAGE T1-weighted sequence (TR = 7.7ms, TE = 3.6ms, FOV = 240 × 240mm^2^, flip angle: 8°, resolution: 1mm^3^, number of slices: 160, slice thickness: 2.2mm, slice orientation: sagittal).

### 2.5. PREPROCESSING OF FMRI DATA

Conventional pre-processing steps for fMRI data were applied to each individual run in native three-dimensional space, using FSL’s Expert Analysis Tool FEAT (v6.00; fsl.fmrib.ox.ac.uk/fsl/fslwiki). The following steps were included: Motion correction using MCFLIRT (Jenkinson, 2002), brain extraction using automated brain extraction tool BET (Smith, 2002), high-pass filtering (100Hz), slice-time correction, and spatial smoothing using a 3mm FWHM (full width at half maximum) Gaussian kernel using FEAT. Functional data was aligned to structural images initially using FLIRT (Jenkinson &Smith, 2001), and optimised using boundary-based registration (Greve &Fischl, 2009). BOLD EPI data was assessed for excessive motion using motion parameter estimates from MCFLIRT. If the functional data from a participant showed greater than 1.5mm (half the voxel size) of absolute mean displacement, this participant was excluded from all further analysis.

To reduce physiological noise artifacts, these CSF and white matter were used to extract scan-wise time series which were then added to the model as nuisance regressors in addition to the standard motion parameters.

Structural images were transformed to Montreal Neurological Institute (MNI) standard space using nonlinear registration (FNIRT), and the resulting warp fields were applied to the functional statistical images.

### 2.6. DEFINITION OF REGIONS OF INTEREST

We used each individual participant T1-weighted image to create a cortical surface reconstruction by means of Freesurfer (Fischl et al., 1999). We identified regions of interest (ROIs), specifically SI, anatomically for each subject based on the probabilistic Brodmann area parcellation provided by Freesurfer (Fischl, 2012). More specifically, an S1 hand ROI was defined by combining Brodmann areas (BA) 1, 2, 3a, and 3b. We then converted this S1 ROI to volumetric space. Any holes were filled and non-zero voxels were mean dilated. Next, the axial slices spanning 2cm medial/lateral to the hand area (Yousry et al., 1997) were identified on the 2mm MNI standard brain (min-max MNI z-coordinates=40-62). This mask was non-linearly transformed to each participant’s native structural space. Finally, we used this mask to restrict the S1 ROI and extracted an S1 hand area ROI. Similar ROI definition has been previously used (Diedrichsen et al., 2013; Ejaz et al., 2015; Wiestler &Diedrichsen, 2013). The S1 hand area ROI was used to both extract time-binned estimates as well as to decode information about the stimulus site during the delay period.

### 2.7. UNIVARIATE ANALYSIS

First-level parameter estimates were computed per run using a voxel-based general linear model (GLM) based on the gamma hemodynamic response function. Time series statistical analysis was carried out using FILM (FMRIB’s Improved Linear Model) with local autocorrelation correction.

To find neural correlates of WM we contrasted beta estimates from the delay period during memory trials to those in no memory trials. We then used a fixed effects higher-level analysis to average activity across runs for each individual participant. Finally, to make inferences on the population level, we computed a mixed effects analysis (Flame 1). From this we obtained statistical group maps (Z-statistic images) for each contrast of interest, e.g. contrasting memory delay activity to no memory delay activity. Z-statistic images were thresholded using clusters determined by Z > 3.1 and statistical significance was determined at the cluster level (p < .05 family-wise-error-corrected (FWE)).

To further explore whether finger specific activity levels were maintained in a somatotopic fashion, we first computed somatotopic ROIs by contrasting finger-specific activity during the first stimulation. We did this by contrasting activity associated with right index stimulations to right little finger stimulations, which elicited a finger specific map (finger cluster) in the lateral part of S1 while the reverse contrast revealed more medially located activity (**Fig. 3B**). These S1 activity maps were in line with previous findings on finger somatotopy.

We then compared z-scored beta estimates between trials where either the index or the little finger was stimulated within each finger ROI. Again, we computed a fixed-effects analysis as mentioned before. We extracted the beta estimates separately for each participant within each pre-defined finger cluster.

**Table 1.** Identified brain regions in which the local activity during the delay period reflected tactile WM processing as shown in Fig. 3. In a next step we parametrically modulated the delay activity, assuming a U-shaped activity during the delay period. For visualization purposes overlap of brain areas was just added the additional z stats for the parametric modulation results. All z-statistic images were thresholded using clusters determined by Z > 3.1 and p < .05 family-wise-error-corrected (FWE) cluster significance. Mean Fischer Z indicates peak z-values. Areas were labeled according to the Juelich Histological Atlas and Havard-Oxford (Sub-)cortcial Structural Atlas (Eickhoff et al., 2005). MFG= medial frontal gyrus, IFG = inferior frontal gyrus, PMC = pre-motor cortex, SMA = supplementary motor area, IPL = inferior parietal lobule, SPL = superior parietal lobule, SMG = supramarginal gyrus.

| Anatomical region | Peak MNI Coordinates |  |  | Mean Fisher Z | U-shaped activity |
| --- | --- | --- | --- | --- | --- |
|  | X | Y | Z |  |  |
| Left Frontal Pole | -42 | 52 | 2 | 4.33 | 3.29 |
| Right Frontal Pole | 40 | 46 | 32 | 3.86 | 3.16 |
| Left IFG | -52 | 16 | 2 | 5.14 | 6.44 |
| Right IFG | 58 | 10 | 18 | 4.59 | 5.47 |
| Left MFG | -44 | 30 | 34 | 4.45 | - |
| Left PMC (+SMA) | -4 | 8 | 50 | 5.49 | 5.81 |
| Right PMC (+SMA) | 8 | 20 | 46 | 5.78 | 5.66 |
| Left S2 | -56 | -23 | 20 | 3.25 | 5.75 |
| Right S2 | 54 | -14 | 16 | - | 5.14 |
| Left S1 | -54 | -22 | 46 | - | 5.63 |
| Right S1 | 56 | -16 | 40 | - | 5.75 |
| Left IPS | -42 | -52 | 48 | 5.14 |  |
| Left IPL | -56 | -20 | 28 | 4.78 | 5.52 |
| Right IPL | 46 | -48 | 52 | 4.86 | 5.67 |
| Left SMG | -42 | -46 | 40 | 4.77 | - |
| Right SMG | 51 | -32 | 46 | 4.16 | - |
| Left SPL | -50 | -48 | 58 | 4.64 | 4.97 |
| Left Insular cortex | -30 | 24 | 6 | 5.61 | 5.41 |
| Right Insular cortex | 36 | 22 | -2 | 5.41 | 5.32 |
| Left Accumbens | -12 | 14 | -6 | 5.27 | - |
| Right Accumbens | 10 | 12 | -4 | 4.16 | - |
| Left Putamen | -18 | 6 | -10 | 4.96 | 4.42 |
| Right Putamen | 20 | 10 | -8 | 4.34 | 3.71 |
| Left Caudate | -12 | 18 | 0 | 4.80 | 5.45 |
| Right Caudate | 14 | 18 | 4 | 4.97 | 4.32 |
| Left Thalamus | -8 | -24 | -16 | 3.86 | 3.81 |
| Right Thalamus | 2 | -20 | -12 | 3.91 | 3.99 |

Information retention in WM is not always reflected by constant delay activity, especially when the duration of the delay period can be somewhat anticipated (Rose et al., 2016). WM delay activity has been shown to decrease until shortly before memory retrieval when the remembered stimulus information is reactivated as suggested by an increase of neural oscillations in the theta band (Rose et al., 2016). We therefore hypothesized that the BOLD activity level would vary in a U-shaped fashion over the delay period. To test this hypothesis, we conducted a parametric modulation analysis. Our parametric modulation regressor was modelled to predict activity in three consecutive time-bins of the delay period in a U-shaped manner (**Fig. 3A**). The length of each time-bin equalled one TR (i.e., 2s). Since we jittered the delay period between 6 and 8 s, we only modelled the first three time-bins of the delay period (2-6s). The remaining time of the delay was modelled as a separate regressor of no interest. By contrasting memory and no memory trials, we obtained Z-statistic images. These images were thresholded using clusters determined by Z > 3.1 and a familywise error–corrected cluster significance threshold of p < 0.05 was applied to the suprathreshold clusters.

To further visualize the results of the parametric modulation, we extracted activity estimates per time-bin. To do so, we modelled each time-bin of the delay period separately in a voxel-based general linear model (GLM) based on the gamma hemodynamic response function. The remaining time of the delay was modelled as a separate regressor of no interest. We then extracted the z-scored estimates per time-bin within the previously defined S1 Hand area ROI. All statistical maps were overlaid onto a MNI152 standard-space T1-weighted average structural template image (**Appx. A.1** and **B.2**) and projected onto a cortical surface using Connectome’s Workbench (Marcus et al., 2011).

### 2.8. VARIANCE INFLATION FACTOR (VIF)

To test whether multicollinearity between the parameter estimates in our GLM was sufficiently low, we calculated the variance inflation factor (VIF). This represents how much the variance of an individual regressor can be explained due to correlation to other regressors in our model (Zuur et al., 2010). For each variable, VIF was computed by the following formula:

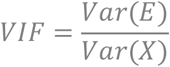

where *Var(E)* reflects the mean estimation variance of all regression weights (stimulation and information storage regressors for each finger) while *Var(X)* reflects the mean estimation variance in case all regressors would be estimated individually. A VIF of 1 indicates total absence of collinearity between the regressor of interest and all other regressors in our GLM while a large VIF signals a serious collinearity problem. There is no clear threshold for acceptable multicollinearity.

Previous literature however recommends that the VIF is ideally smaller than 2.5 (Johnston et al., 2018). In our case, the VIF was 1.45, averaging across regressors reflecting te first stimulation, the delay and the second stimulation. We additionally computed the VIF for the regressors relating to the stimulation periods and the time-binned delay periods (i.e. stimulation one, delay (0-2s), delay (2-4s), delay (4-6s) and stimulation two). As expected, the resulting VIF was on average 11.38, signaling concerning multicollinearity. Since the WM related activity within the time-bins is correlated, this result was expected, but also means our findings of our time-binned analysis should be interpreted with caution.

### 2.9. MULTIVARIATE PATTERN ANALYSIS

#### 2.9.1. MVPA

We used multi-voxel pattern analysis (MVPA) to decode which finger was stimulated based on activity during the delay period. This analysis was conducted for voxels within the S1 hand area mask that have been shown to possess fine-grained finger representations. First-level parameter estimates were computed for all events of each trial and each participant using a voxel-based general linear model (GLM) in SPM (v12) based on the gamma hemodynamic response function. This resulted in 192 beta estimates (48 trials x 4 runs) during the delay period per participant across both conditions. 96 beta estimates for each memory and no memory condition.

We trained a linear classifier (support vector machine, SVM) to predict which finger was stimulated in a specific trail based on the respective delay period activity using the nilearn toolbox (Abraham et al., 2014). We calculated classification accuracies using a leave-one-run-out cross-validation approach. The accuracies were averaged across folds, resulting in one accuracy per condition and per participant. To approximate the true chance level, we shuffled the condition labels using 1000 permutations (Ojala &Garriga, 2009), as implemented in the scikitlearn toolbox (Pedregosa, 2011). Then, we calculated permutation-based p-values based on the following formula:

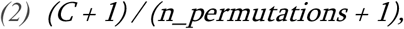

where *C* is the number of permutation scores that were higher than the true accuracy. To assess the statistical significance of each ROI’s decoding accuracy, we used one sample t-tests against the approximated chance level (Stelzer et al., 2013). We further used a paired t-tests to compare classification accuracies between the WM and non-WM conditions.

#### 2.9.2. REPRESENTATIONAL SIMILARITY ANALYSIS (RSA)

RSA has the ability to identify the invariant representational structure of fingers independent of amplitude, shape and exact location of activated brain regions during the WM task (Ejaz et al., 2015). It allowed us to obtain a measure of how distinguishable somatotopic representations between working-memory and no working-memory trials are. We computed representational distances between activity patterns related to different fingers (index vs.little finger) for both working-memory and no working-memory conditions. The distances were obtained using a prewhitened crossvalidated Malhalnobis distances (Crossnobis distances, (Diedrichsen et al., 2013; Walther et al., 2016). We obtained voxel-wise parameter estimates (betas) for each finger * memory condition * timepoint versus rest (using univariate analysis) and residuals of our GLM within the hand area of S1. These betas were prewhitened using the residuals. Based on the prewhitened betas, we computed squared Mahalanobis distances between all possible finger*condition*timepoint combinations for each fold (i.e. run) and averaged them across folds. A distance greater than 0 reflects dissociable cortical representations while 0 shows no dissociation. The distance measures between all possible representations were assembled in a representational dissimilarity matrix (RDM). For visualization, we only extracted distances between the finger representations during memory vs. no memory for each time point (f1, delay and f2) of the task.

### 2.10. STATISTICAL DATA ANALYSIS

To detect outliers, we used the robustbase toolbox (Finger, 2010). **Sn** identifies an outlier (xi) if the median distance of xi from all other points, was greater than the outlier criterion (λ=3) times the median absolute distance of every point from every other point:

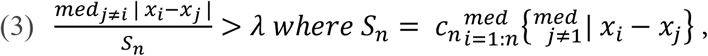

where cn is a bias correction factor for finite sample sizes (Rousseeuw &Croux, 1993). We detected no outliers for behavioral data that had to be excluded from any further analysis.

Before conducting any repeated measures ANOVA testing, we validated the assumptions for normaility and spherecity using a Shapiro-Wilk and Mauchy test. Effect sizes of different variables were measured using eta squared. ANOVA analysis was done using the pingouin toolbox (Vallat, 2018). P-values were Greenhouse-Geisser corrected if sphericity could not be assumed. T-statistics were corrected for multiple comparisons using one-step Bonferroni correction.

Bayesian analysis was carried out using pingouin toolbox for the main comparisons to investigate support for the null hypothesis. Following the conventional cut-offs, a BF smaller than 1/3 is considered substantial evidence in favor of the null hypothesis. A BF greater than 3 is considered substantial evidence, and a BF greater than 10 is considered strong evidence in favor of the alternative hypothesis. A BF between 1/3 and 3 is considered weak or anecdotal evidence (Dienes, 2014; Kass &Raftery, 1995).

## 3. RESULTS

### 3.1 BEHAVIORAL PERFORMANCE WAS BETTER WITH GREATER FREQUENCY DIFFERENCES

A two-way ANOVA showed that behavioral performances differed significantly depending on the frequency differences between the first and second vibrotactile stimulus (F(2, 156) = 5.06, p < .001, η^2^ = 0.06); **Fig. 2A**), but not between stimulated fingers (F(1, 156) = .44, p = .51, η^2^ < 0.01). There was no interaction effect; F(1, 156) = .88, p = .42, η^2^ = 0.01). A post hoc test (Tuckey’s HSD) on pairwise comparisons on frequency differences pairs revealed that discrimination accuracy was significantly different between 2 and 6 Hz differences (*q =* 4.48, *p* < .01) and showed no significant difference for the rest of the pairs (4 Hz vs 6 Hz: *q =* 2.67, *p* = .15 and 2 Hz vs 4 Hz *q =* 1.81, *p* = .41). We further analyzed whether behavioral performance changed across runs despite our efforts to re-adjust the detection threshold (**Fig. 2B)** and found only insignificant differences across runs (F(3, 2496) = 2.0, p = .11, η^2^ < 0.01) and across fingers (F(1, 2496) = .48, p =.49, η^2^ < 0.01), and no significant interaction effect (F(3, 2496) = .47, p = .47, η^2^ < 0.01).

**Fig. 2.**
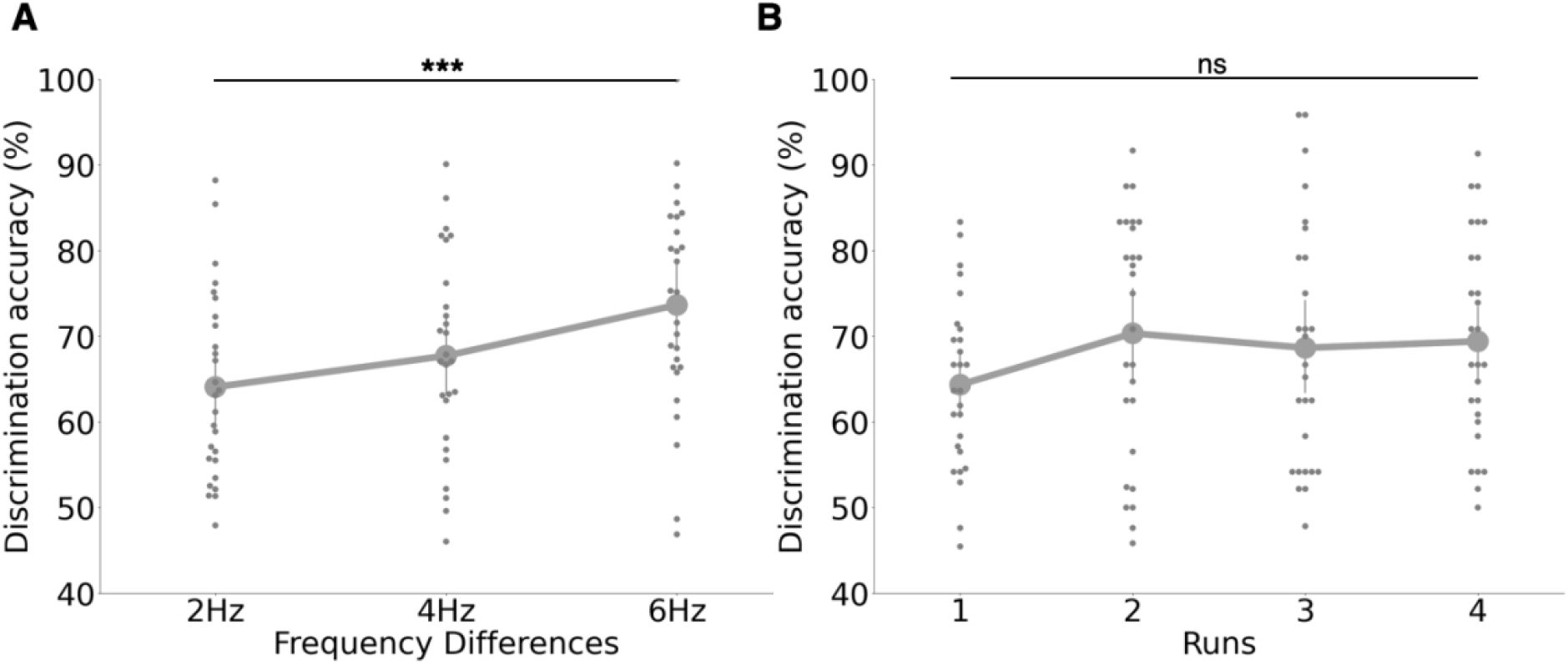
Behavioral performance results. **A**. Discrimination accuracy (% of correct answers) improved when frequency differences were larger. The blue dots reflect the group mean and the blue error bars indicate the standard deviation of the discrimination accuracy per frequency difference across the whole experiment. **B**. Behavioral performance was not significantly different across runs. The blue dots reflect the group mean and the blue error bars indicate the standard deviation of discrimination accuracies across each run. This demonstrates that our SDT criterion assured stable discrimination accuracies across runs. Grey dots represent individual participants’ results. *** = p < .001; ns = non-significant.

### 3.2. VIBROTACTILE WORKING MEMORY RECRUITS A DISTRIBUTED BRAIN NETWORK

We first determined brain areas that were more activate during the delay period in the memory compared to the no memory condition. We found that WM processing involved a distributed brain network: i.e., bilateral frontal lobe, bilateral (medial/inferior) frontal gyrus (MFG, IFG), bilateral pre-motor cortex and supplementary motor area (PMC, SMA), contralateral secondary somatosensory cortex (S2), bilateral inferior parietal lobule (IPL), bilateral superior parietal lobule (SPL), bilateral supramarginal gyrus (SMG), bilateral caudate, bilateral thalamus, bilateral nucleus accumbens, and bilateral insula (**Fig. 3A, Table 1** and **Appx. A.1**). As in previous human fMRI studies, this univariate analysis did not reveal significant S1 activity.

**Fig. 3.**
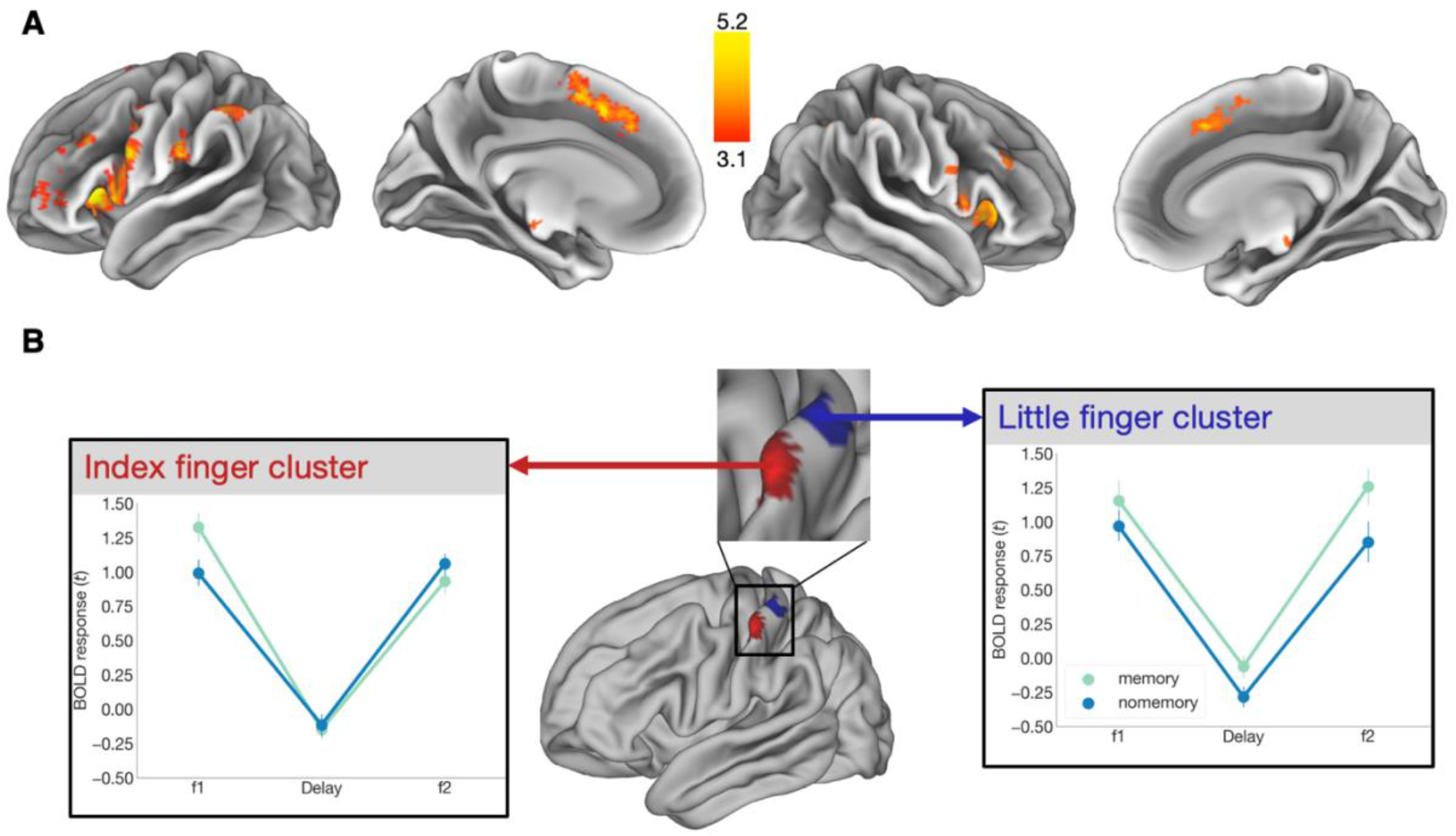
Univariate group results. **A**. We determined brain regions that were more activate during the delay period of the memory compared to the no memory condition. A statistical map (Z > 3.1) was obtained by contrasting delay period activity in memory trials to no memory trials. The map was separately projected onto a cortical surface contralateral (*top*) and ipsilateral (*bottom*) to the stimulus site. WM related activity resided in a network of brain regions (for more details, see **Table 1**). **B**. S1 areas activated during index (red) and little (blue) finger stimulation (*middle*). Clusters activated during index and little finger stimulation Z-statistic images were thresholded using clusters determined by Z > 3.1, p < .05 family-wise-error-corrected (FWE) cluster significance and were projected onto a cortical surface. Finger maps were located in contralateral S1. We then extracted the t-scored beta estimates within these finger-specific S1 areas during memory and no memory trials at different timepoints (f1, delay and f2) by contrasting cluster-specific finger activities (e.g. index finger cluster: index finger memory trials > little finger memory trials; bottom). Point plots are centered at the mean and error bars reflect the standard error.

Furthermore, we contrasted activity levels during vibrotactile stimulation between fingers (i.e., index>little and little>index finger), and, as expected, observed separated finger representations in contralateral S1(**Fig. 3B**, *middle*) with the little finger being represented more medially than the index finger (Besle et al., 2013; Kikkert et al., 2021; Kolasinski et al., 2016; Martuzzi et al., 2014; Sanchez Panchuelo et al., 2018; Sanders et al., 2019). Finally, we assessed whether finger-specific activity (e.g. index finger related activity which was greater within the index finger cluster) was modulated within these finger-specific clusters during memory compared to no memory trials at different timepoints of the task (f1, delay and f2). Our repeated measures ANOVA revealed that t-scored beta estimates obtained from index finger cluster did significantly differ between time points (timepoints main effect: F(1, 52) = 52.15, p_*corr*_ < .001, η^2^ = .49), but not between conditions (conditions main effect: F(2, 26) = .14, p_*corr*_ = .5, η^2^ < .05) and. We also found no significant interaction effect (F(2, 52) = 2.04, p_*corr*_ = .14, η^2^ < .05; **Fig. 3B**, *left*).

Extracted t-scored beta estimates from the little finger cluster revealed a time point main effect (F(1, 52) = 38.63, p_*corr*_ < .001, η^2^ = .45) and a condition main effect F(1, 26) = 8.75, p_*corr*_ < .01, η^2^ < .05), but no interaction effect (F(2, 52) = .54, p_*corr*_ = .58, η^2^ < 0.05; **Fig. 3B**, *right*).

### 3.3. TEMPORAL MODULATION OF DELAY PERIOD ACTIVITY IN CONTRALATERAL S1

We then examined how brain activity temporally changed during the delay period. We parametrically modulated the delay period regressors by the hypothesized U-shaped activity changes and computed the associated statistical maps. A U-shaped activity modulation was found in a similar network of brain regions as displayed in **Fig. 4** and **Appx. B.1**. In addition, we also found significant changes in BOLD signal in bilateral S1 and S2. S1 activity overlapped with the area that usually represents the hand.

**Fig. 4.**
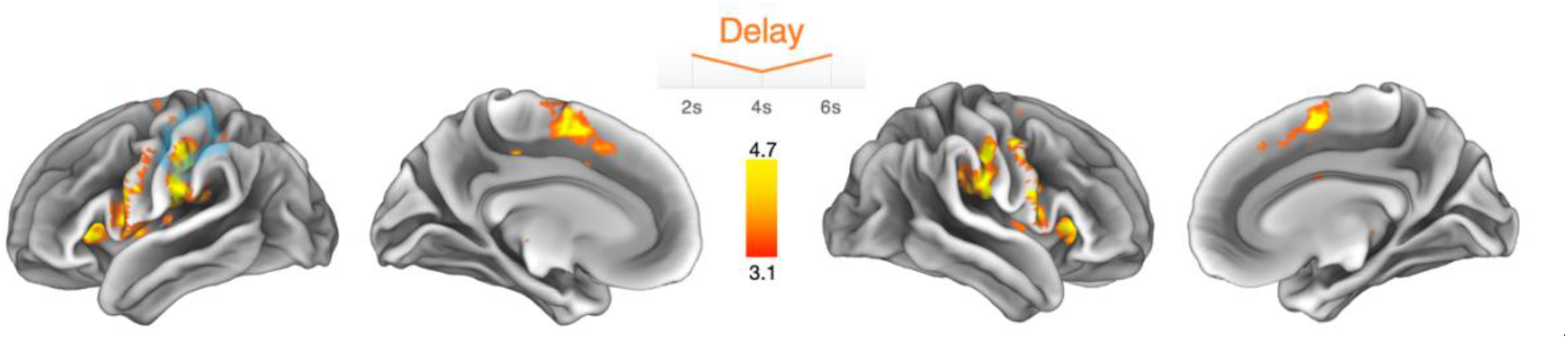
U-shaped parametric modulation of WM-related activity. **A**. Brain regions exhibiting u-shaped modulated delay activity patterns (see insert at the top reflecting the parametric modulator entered into the GLM) during the delay period (2-6s). The contrast shows the difference between the memory and no memory condition in the contralateral (*left*) and ipsilateral hemisphere (*right*). The area highlighted in blue represents the S1 hand area.

### FINGER-SPECIFIC S1 WM REPRESENTATIONS

We hypothesized that executing a WM task would modulate finger specific representations in S1. To test this hypothesis, we used MVPA to decode the stimulated finger (i. e., index versus little finger) during the first stimulation (f1), during the delay period, and the second stimulation (f2) separately for memory and no memory trials (**Fig. 5A**). We did this in the contralateral S1 ROI, which has shown to possess fine-grained finger representations (Besle et al., 2013; Martuzzi et al., 2014; Sanchez Panchuelo et al., 2018; Sanders et al., 2019), and a control white matter ROI.

**Fig. 5.**
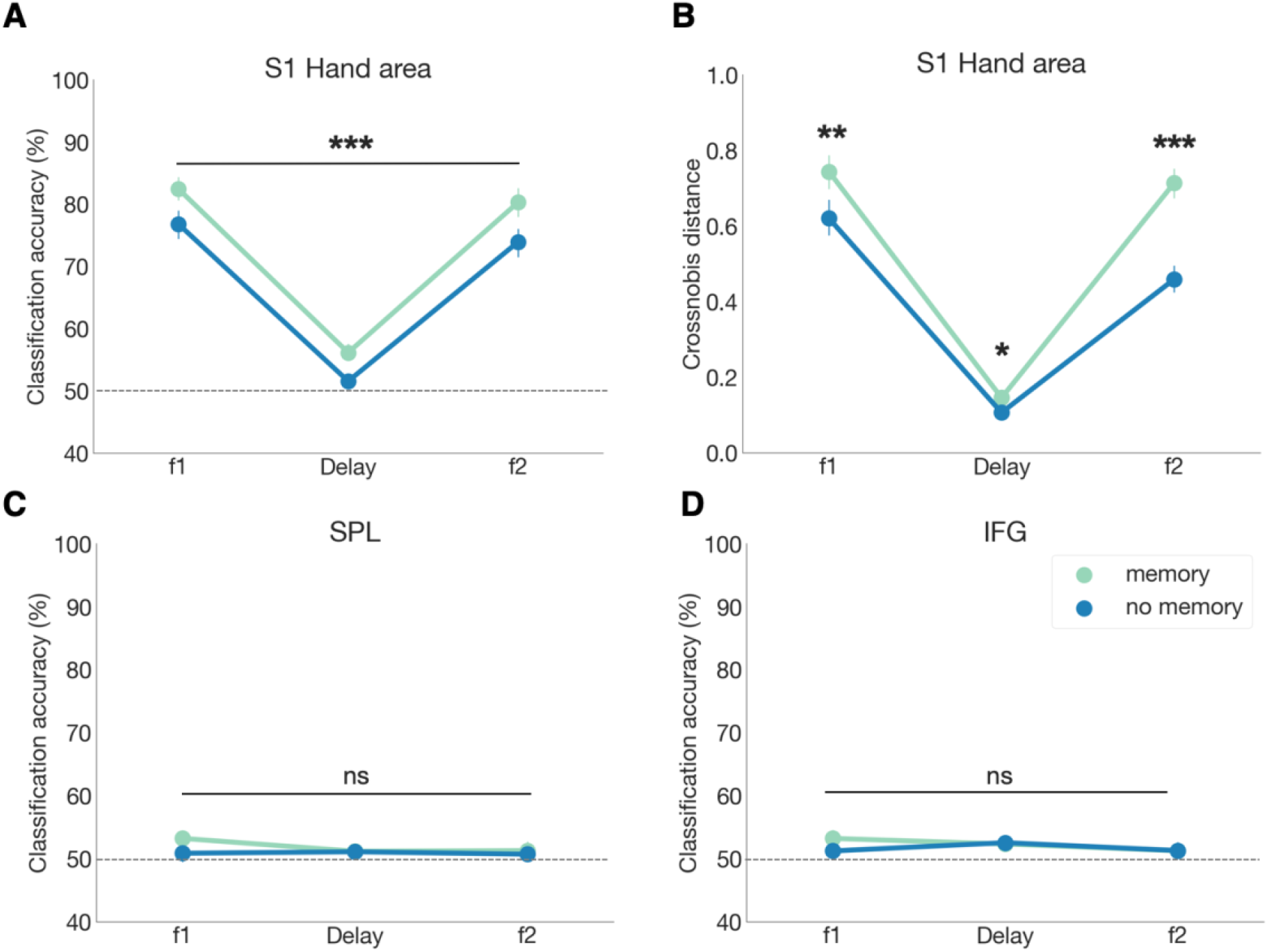
Multivariate results on somatotopic modulations. We investigated whether activity patterns within a ROI with fine-grained finger somatotopy became more distinct at different timepoints (f1, delay and f2) during memory trials compared to no memory trials. **A**. Classification accuracies (index vs. little finger) based on activity patterns in hand area within contralateral S1. **B**. Cross-validated Mahalanobis (Crossnobis) distances between fingers x condition in the same ROI. **C**,**D**. Further explorative analysis of ROIs that previously were indicated to be involved in vibrotactile WM, but where no fine-grained somatotopy is assumed. The point plots are centered at the mean and error bars reflect the standard error. Grey dotted lines reflect the theoretical chance level. If interaction effects were significant, pairwise comparisons results for comparing memory vs. no memory conditions separately for each time point are indicated by * p < .05, ** p < .01 *** p < .001. S1 = primary somatosensory cortex, IFG = inferior frontal gyrus, SPL = superior parietal lobule.

Our repeated measures ANOVA revealed that classification accuracies obtained from contralateral S1 hand area significantly differed between time points (f1,delay and f2; time point main effect: F(2, 52) = 156.52, p_*corr*_ < .001, η^2^ = .77) and between memory and no memory trials (condition main effect: F(1, 26) = 48.42, p_*corr*_ < .001, η^2^ < .05). However, we found no interaction effect (i.e., F(2, 52) = .5, p_*corr*_ = .59, η^2^ < 0.001).

Moreover, we showed that differences between memory versus no memory trials reached significance. This was confirmed by pairwise comparisons (t(26) = 6.39, p_*corr*_ < .001; BF10 = 1.95e^04^ with the Bayes factor (BF) showing extreme evidence in favor of the WM-related modulation.

Permutation tests allowed us to obtain approximated chance levels. We observed that classification accuracies were significantly greater than this chance level for f1 and f2, irrespective of whether it was a memory or no memory trial (*t(25,25)* >= 13.24, *p* < .001). During the delay period, however, classification exceeded the chance level only for memory trials (*t(26)* = 4.55, *p* < .001) but not for no memory trials (*t(26)* = 1.5, *p* = .15).

Furthermore, we also investigated the representational geometry of memorized tactile stimuli (**Fig. 5B**). We hypothesized that executing a WM task would also modulate the finger representational geometry in S1. Our repeated measures ANOVA revealed that the cross-validated Mahalanobis (Crossnobis) distances obtained from contralateral S1 hand area significantly differed between time points (f1,delay and f2; time point main effect: F(2, 52) = 151.21, p_*corr*_ < .001, η^2^ = .73) and between memory and no memory trials (condition main effect: F(1, 26) = 39.81, p_*corr*_ < .001, η^2^ = .06). We also found an interaction effect (i.e., F(2, 52) = 28.65, p_*corr*_ < .001, η_2_ < 0.05). We again could show that geometrical differences of finger representations between memory versus no memory trials reached significance when independently tested for each of the three time points and, particularly, for the delay period (i.e. in the absence of any tactile stimulation). This was confirmed by pairwise comparisons (f1 memory vs. f1 no memory: t(26) = 4.09, p_*corr*_ < .01; BF10 = 81.73 , Delay memory vs. delay no memory: t(26) = 2.9, p_*corr*_ < .05; BF10 = 5.96, f2 memory vs. f2 no memory: t(26) = 7.4, p_*corr*_ < .001; BF10 > 100, with the Bayes factor (BF) showing substantial evidence in favor of the null hypothesis.

We also explored further ROIs located on superior parietal lobe (SPL; **Fig. 5C**) and inferior frontal gyrus (IFG; **Fig. 5D**) that have been implicated to be involved in vibrotactile WM (Schmidt et al., 2017, 2021; Schmidt &Blankenburg, 2018). ROIs were based on clusters obtained in univariate analysis (DelayMemory > DelaynoMemory). We found no significant differences in classification accuracies based on activity patterns within contralateral IFG (time point main effect, F(2, 52) = 0.93, p_*corr*_ = 0.4, η^2^ < .05; condition main effect, F(1, 26) = 0.38, p_*corr*_ = 0.55, η^2^ < .05) and time point x condition interaction effect F(2, 52) = 1.28, p_*corr*_ = .28, η^2^ < .05) as well as within contralateral SPL (time point main effect, F(2, 52) = .75, p_*corr*_ = .46, η^2^ < .05; condition main effect, F(1, 26) = .85, p_*corr*_ = .37, η^2^ < .05 and time point x condition interaction effect F(2, 52) = .68, p_*corr*_ = .51, η^2^ < .05). Together, these results suggest that during a vibrotactile WM task finger representations are likely modulated in S1 and not in other areas of the WM network, even in the absence of any tactile stimulation.

## 4. DISCUSSION

Tactile decision making based on mentally represented information is an essential part of higher cognitive processes. In the absence of sensory stimuli, decisions are made with information stored in WM (Christophel et al., 2017). A more profound insight into how the brain stores information in WM is essential to understand human cognition. This includes not only the involvement of sensory regions in WM but also how their topographic organization might be modulated. In the present study, we demonstrated that finger-specific, somatosensory information represented in S1 is modulated by cognitive processes relevant for performing a vibrotactile frequency discrimination task. Our main finding was that S1 activity during the delay period (i.e. in the absence of any tactile stimulation) contained information on which finger received somatosensory input even though the average BOLD activity in S1 did not differ between conditions. Importantly, when participants received the same stimuli in a no memory condition, classification accuracies and representational dissimilarities were significantly lower indicting that our main finding was specific to the cognitive task.

We propose that keeping task-relevant tactile information in WM modulated finger representations in S1 due to top-down control mechanisms that sharpens tuning curves of neurons in S1 and might be associated with attentional control.

### 4.1. SOMATOTOPIC MODULATION IN S1 IS MAINTAINED DURING WM DELAY PERIOD

When exposed to a stimulus, stimulus-selective neurons are activated which is can be quantified as a tuning curve. Previously, specific neurons have shown ‘tuned’ responses to different features of the stimulus, e.g. stimulus location or stimulus orientation (Campbell et al., 1968; Henry et al., 1974; Scobey &Gabor, 1989). For instance, responses of neuron populations in primary visual cortex (V1) can be modulated by changes of stimulus orientation (Hubel &Wiesel, 1968). The product of multiple tuning curves can be defined as the neural population code (Ben-Yishai et al., 1995; Georgopoulos et al., 1986). A linear decoder has the ability to capture the information kept in the neural population code (Kriegeskorte &Wei, 2021). Similarly, neural tuning also determines the representational geometry in the multivariate response space and these changes in geometry can be detected by RSA (Kriegeskorte &Wei, 2021).

Even in the absence of sensory input, the somatosensory cortex in non-human primates has shown to possess neurons whose tuning curves properties depend on the frequency of the a priori presented vibrotactile stimulus (Romo &Salinas, 2003). Neuroimaging studies in humans using MVPA have further revealed that tactospatial information is retained during the delay period by a neural population code in S1, SPL, PMC and posterior parietal cortex (PPC) while preserved frequency information is reflected by activity patterns in dorsal PMC, SMA and IFG (Schmidt et al., 2017; Schmidt &Blankenburg, 2018). Note that we did not attempt to directly decode task-relevant features of the tactile WM content. However, it is likely that immediate modulation of the somatotopic map in S1 is inherent to performing the WM tasks since it involves encoding stimulus location information. This could explain why previously such modulation was also observed during (i) attempted finger movements that activated S1 in an effector specific manner even in the absence of any motor activity and self-generated sensory feedback (Kikkert et al., 2016, 2021; Wesselink et al., 2019), (ii) viewing touches (Kuehn et al., 2018), actively imagining touch (Schmidt &Blankenburg, 2019), (iii) or attending to individual finger stimulation (Puckett et al., 2017). Likewise, keeping tactile stimuli in WM (as in our case) appears to be sufficient to generate a more dissociable somatotopic map in S1, a process that might aid tactile decision making. The advantage of the topographic arrangement in S1 might be a reduction in time of information transmission and thus metabolic expensive connections (for review,(Manni &Petrosini, 2004). When afferent input from adjacent body parts enters the brain, their cortical representations might facilitate discrimination between sensory information from different spatial locations.

Even though our results did not allow to draw any conclusions on its task-relevance, there exists evidence that tactile decision making depends on neural activity in S1. Two transcranial magnetic stimulation (TMS) studies in humans found impaired behavioral vibrotactile frequency discrimination performance when TMS pulses were applied to either contralateral S1 during the beginning (Harris et al., 2002) or to ipsilateral S1 at the end of the delay period (Zhao &Ku, 2018). This suggests that when performing a vibrotactile WM task neural activity in S1 appears to be task-relevant and not only restricted to the contralateral hemisphere.

It is uncertain which specific cognitive mechanism might have driven the observed modulation. A discrimination task usually requires enhancement of neural activity to relevant stimuli and suppress activity to irrelevant stimuli. Such amplification of relevant information has been conceptualized as generalized models of attention (for review, Burton &Sinclair, 2000). Attention has been suggested as an integral part of performing WM tasks (Cowan et al., 2013; Logie &Cowan, 2015; Schmidt &Blankenburg, 2018, 2019) as it filters information by sensory modality or by body location (Gomez-Ramirez et al., 2011) and, thereby, attention regulates what will be cortically represented and what will not (Desimone &Duncan, 1995). Furthermore, it has been shown that volitionally directing attention towards a spot on the body surface which was tacitly stimulated increases BOLD responses in S1 (Nelson et al., 2004; Puckett et al., 2017; Sterr et al., 2007). Even the expectation over being stimulated on a specific finger was sufficient to modulate neural activity in S1 in a somatotopic manner (Roland, 1981, Drevet et al. 1995). Both our univariate and multivariate results support this notion. During both time points of stimulation (f1 and f2) we observed significantly greater finger-specific BOLD responses in S1 and obtained higher classification accuracies and crossnobis distances for the memory than the no memory condition.

There is accumulating evidence that attention modulates tuning curves in the specific sensory modality (Bisley, 2011; McAdams &Maunsell, 1999; Reynolds et al., 2000). By sharpening tuning curves attention contributes to increased discrimination performance ((Bartsch et al., 2017), but see also Seriès et al., 2004). According to the sharpening hypothesis, predicted or attended stimuli lead to suppressed neural activity, while increasing the information content (Murray &Wojciulik, 2004). This might explain why we observed no difference in BOLD responses in S1 during the delay period of memory trials compared to no memory trials, while we found significantly higher classification accuracies and representational dissimilarities between fingers that indicated the presence of greater amounts of information.

In summary, our findings are in line with the idea that greater representational dissimilarities during WM might reflect attentional top-down control which optimally tunes somatotopic finger representation while vibrotactile stimulation is received (i.e. during f1 and f2) but, more importantly, also during the delay period.

### 4.2. TEMPORAL MODULATION OF BRAIN ACTIVITY DURING THE WM DELAY PERIOD

In line with previous research (Pasternak &Greenlee, 2005; Preuschhof et al., 2006), we identified a widespread parieto-fronto-insular network which is typically involved in tactile WM. We could show that fronto-partial areas, insular cortices, subcortical regions (i.e. caudate and thalamus), S1 and S2 exhibited a U-shaped activity profile across the delay period, which might provide a glimpse into the temporal modulation of WM related brain activity (Cohen et al. 1997). Neurophysiological models of WM have proposed that vibrotactile information is processed in S1, relayed to S2 (Kalberlah et al., 2013), which possesses a reciprocal connection with S1 and disseminates information via insular projections to frontal and parietal regions (Wang et al., 2013). Task-relevant modulation could occur through prefrontal cortex (PFC) by gating the flow of vibrotactile information to S1. Indeed, the prefrontal-thalamic inhibitory system could play a key role in boosting relevant stimulus information while reducing irrelevant ones (Staines et al., 2002). It is likely that the final increase in delay activity might be driven by anticipation of increasing attentional demands, although we jittered the length of the delay period to prevent for anticipatory activity (Rose et al., 2016). Therefore, it is very likely that attentional mechanisms are driving our results. However, as mentioned before, WM and attention are thought to be closely intertwined (LaBar et al., 1999; Naghavi &Nyberg, 2005) and our set-up did not allow to disentangle these aspects.

### 4.2. INTERPRETATIONAL ISSUES

Our multivariate analysis results suggest that a WM task modulates representational tuning in S1. However, our memory delay period was relatively short (6-8s) compared to other fMRI studies investigating vibrotactile WM that often employ delays of 12s to avoid any carry-over effects (Schmidt &Blankenburg, 2018; Wu et al., 2018). Nonetheless, it seems unlikely that the WM related somatotopic activity patterns observed in S1 were merely a residue of stimulus evoked activity due to the sluggishness of the BOLD response since we directly compared the memory and no memory condition. If we merely decoded a residue of the tactile stimulation per se, then we would expect similar decoding results for both conditions during the delay period. Indeed, finger stimulations were identical for the memory and no memory conditions. The only difference between these conditions was the cognitive strategy participants employed. While classification accuracies and Crossnobis distances were significantly higher during the memory condition, this was not the case during the no memory condition. Finally, the low VIF of the regressors suggested that activity related to stimulus perception and WM storage could be disentangled by our model.

## 5. CONCLUSIONS

Our results extend previous findings on the involvement of S1 in vibrotactile frequency discrimination. We provide new evidence for the importance of somatotopy during vibrotactile WM, particularly during the delay period, i.e. in the absence of any stimulation. This is remarkable, since the cortical representation of the stimulus location was not essential for solving the WM task suggesting that some aspects of the tactile WM content are represented in a somatotopic fashion. We speculate that finger representations in S1 are modulated during WM through top-down attentional mechanisms which adds to a broader understanding of human cognition.

## 6. ACHKNOLEDGMENTS

We thank Christian Ruff for stimulating discussions. We also want to thank Daniel Woolley and Roger Lüchinger for technical support as well as Alain Plüss and Bianca Badii for help with data collection.

## 8. SUPPLEMENTARY MATERIAL

**Appx. A.1.**
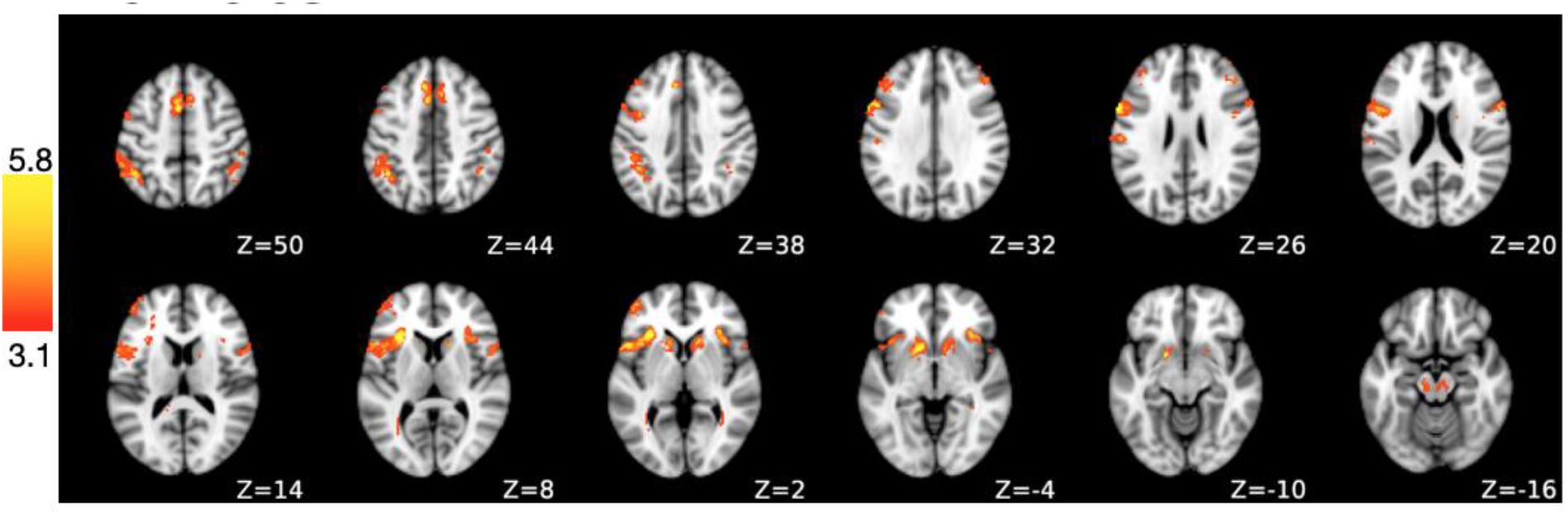
Slice view of univariate group results. We identified brain regions that were more activate during the delay period of the memory compared to the no memory condition. A statistical map (Z > 3.1) was obtained by contrasting modulated delay period activity in memory trials to no memory trials.

**Appx. B.2.**
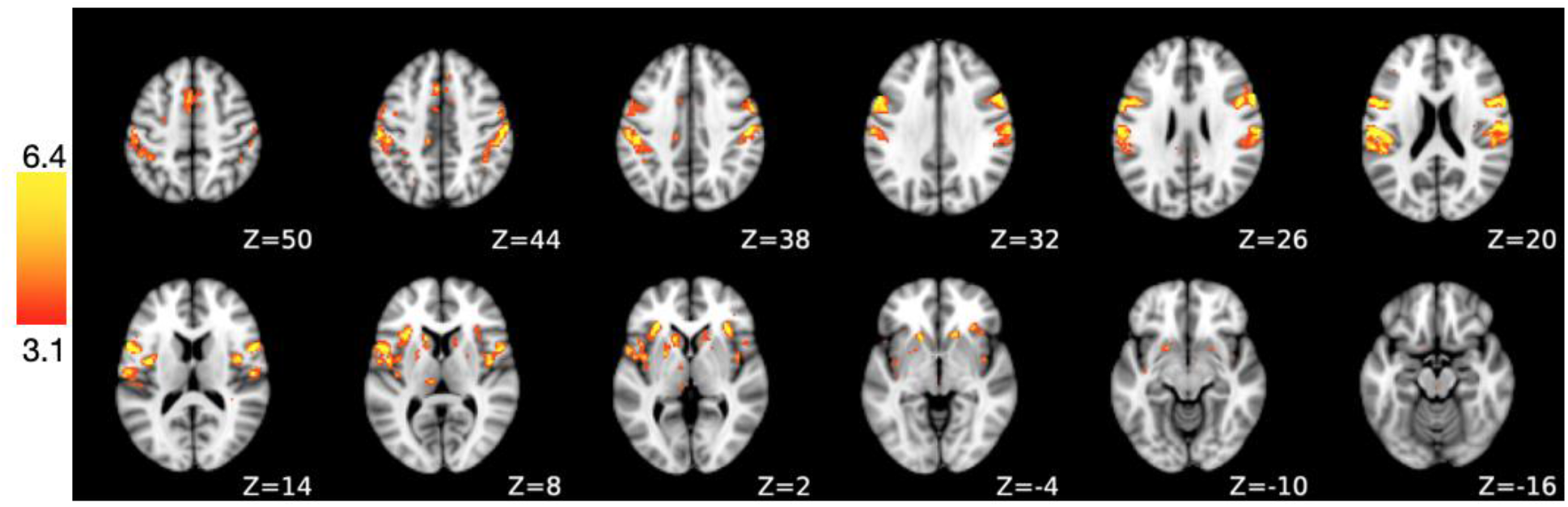
Slice view of parametric modulation results. We identified brain regions exhibiting u-shaped modulated delay activity patterns during the delay period (2-6s). A statistical map (Z > 3.1) was obtained by contrasting delay period activity in memory trials to no memory trials.

